# Analyzing the weak dimerization of a cellulose binding module by sedimentation velocity experiments

**DOI:** 10.1101/638320

**Authors:** Dmitrii Fedorov, Piotr Batys, Maria Sammalkorpi, Markus B. Linder

## Abstract

Cellulose binding modules (CBMs) are found widely in different proteins that act on cellulose. Because they allow a very easy way of binding recombinant proteins to cellulose, they have become widespread in many biotechnological applications involving cellulose. One commonly used variant is the CBM_CipA_ from *Clostridium thermocellum*. Here we studied the dimerization of CBM_CipA_, because we were interested if its solution behavior could have an impact on its use in biotechnical applications. As the principal approach, we used sedimentation velocity analytical ultracentrifugation. To enhance our understanding of the possible interactions, we used molecular dynamics simulations. By analysis of the sedimentation velocity data using a discrete model genetic algorithm we found that the CBM_CipA_ shows a weak dimerization interaction with a dissociation constant *K*_*D*_ of about 87 μM. As the *K*_*D*_ of CBM_CipA_ binding to cellulose is about 0.6 μM, we conclude that the dimerization is unlikely to affect cellulose binding. However, at the high concentrations used in some applications of the CMB_CipA_, its dimerization is likely to have an effect on its solution behavior. The work shows that analytical ultracentrifugation is a very efficient tool to analyze this type of weak interactions. Moreover, we provide here a protocol for data analysis in the program Ultrascan for determining dissociation constants by sedimentation velocity experiments.

## Introduction

Cellulose binding modules (CBMs) are proteins that are found as parts of many different enzymes and other proteins that in one way or the other interact with cellulose. Their role is to bind to cellulose and typically they do so by surface exposed aromatic side chains. It is well known that CBMs play a crucial role in cellulose degradation and enzymatic activity (1). There are up to 40 different families of functionally related, but structurally different proteins that bind to carbohydrates. They form an example of convergent evolution with different protein scaffolds achieving a similar binding function (2). The details of these binding mechanisms differ between the individual families, although they share some common elements such as aromatic-pyranose ring stacking as part of the binding interface.

CBMs have been utilized in a large number of biotechnical applications because they provide an easy way to control the binding of proteins to cellulose. Especially the ability to use recombinant DNA technology to design new structurally engineered proteins with CBMs attached has led to a wide range of applications. For example, the activity of enzymes can be enhanced by increasing their binding to cellulose by CBMs. In one study, different types of CBMs were used to either enhance or inhibit the activity of lytic polysaccharides monooxygenases (LPMO) by swapping between different types of CBMs (3). The activity of hydrolases have also been increased by fusing enzymes to CBMs (4). Immobilization to cellulose is also an application that has been explored. CBMs can be used as a high-capacity purification tag, a targeting molecule or an affinity tag for enzyme immobilization and processing (5). In addition, CBMs can be used to increase a rate of catalysis by creating of association between biocatalysts and substrates (6,7). Other applications include; cell immobilization (8), diagnostics and even protective devices against nerve gas (9).

Because CBMs attach to cellulose surfaces, they can also be used to modify the materials properties of cellulose based materials. An example of this was the improvement of the mechanical properties of paper. A protein constructed to contain two CBMs at each end of a linker, resulted in paper with higher folding endurance (10). In another example, a recombinant silk-protein based adhesive for cellulose included CBMs (11).

Here we focus on the CBM_CipA_ from the *Cellulomonas fimi* cellulosome anchoring protein CipA. The CBM_CipA_ belongs to family III according to the Cazy-classification (2). CBM_CipA_ has a nine-stranded β sandwich like structure with a jellyroll topology and belongs to surface binding type of CBMs (type A). It has a face that contains planar linear strip of aromatic and polar residues, which participates in the interaction with crystalline cellulose (12).

CBM_CipA_ shows a very strong binding to crystalline cellulose, with a dissociation constant (*K*_*D*_) of about 0.6 μM (13–15). This is an order of magnitude stronger than the *K*_*D*_ of, for example, the fungal family I type of CBMs (16). Due to the high affinity of CBM_CipA_ to cellulose, it has a high practical value in applications. Because of its wide use and because any multimerization behavior possibly can influence the use of the protein in the applications, we decided to investigate the multimerization characteristics of CBM_CipA_ in more detail. Furthermore, a possibility of dimerization was indicated in our previous work using CBM_CipA_ as a fusion partner in silk-material forming proteins (11,17,18). For CBM_CipA_, no detailed information of dimerization is available, but the formation of dimers has been reported for the closely related CBD_Cex_ from the enzyme Cex, a β-1,4-glycanase from *C.fimi* (19). As a method to study the solution interactions of the CBM_CipA_ we chose analytical ultracentrifugation (AUC) which is one of the most powerful techniques available for quantifying weak interactions (20). Experiments performed by AUC are either based on analyzing sedimentation velocity or sedimentation equilibrium. When performing sedimentation velocity experiments, one obtains the dissociation constant *K*_*D*_, the diffusion coefficient, as well as information on the anisotropy of the protein in solution (21).

## Materials and Methods

### Protein expression

CBM_CipA_ was codon optimized for *E. coli* and synthetized by GenScript USA (NJ, USA), and cloned into the vector pET28a(+) as described earlier (15). The synthetized genes contained *Bsa*I restriction sites in their 5’ and 3’ ends. The multiple cloning site of the pET28a(+) expression vector was modified to include two *Bsa*I restriction sites by inserting an adapter constructed from oligonucleotides 5’-CATGGGGAGACCGCGGATCCGAATTCGGGTCTC-3’ and 5’-TCGAGAGACCCGAATTCGGATCCGCGGTCTCCC-3’ into *Nco*I and *Xho*I restriction sites. Plasmids were transformed to *E. coli* XL1-blue strain and plated on kanamycin (50µg/mL) containing LB-plates. The CBM_CipA_ contained a 6-His-tag in the C-terminal end. The sequence of the CBM_CipA_ was:

MGNLKVEFYNSNPSDTTNSINPQFKVTNTGSSAIDLSKLTLRYYYTVDGQKDQTFWCDHAAIIGSNGSYNGITSNVKGTFVKMSSSTNNADTYLEISFTGGTLEPGAHVQIQGRFAKNDWSNYTQSNDYSFKSASQFVEWDQVTAYLNGVLVWGKEPLEHHHHHH

*E.coli* strain BL21 (DE3) (ThermoFisher Scientific) was used for CBM_CipA_ production. CBM_CipA_ was expressed using MagicMedia™ (ThermoFisher Scientific) expression medium in accordance with the manufacturers protocol. The purification process consisted of two parts, precipitation by heating and by immobilized metal affinity chromatography (IMAC). The sample was heated up to 70 °C for 30 minutes to precipitate impurities, and then clarified by centrifugation at 5000 rpm. An ÄKTA-Pure liquid chromatography system (GE Healthcare Life Sciences) with a 5mL HisTrap column was used for IMAC purification of the protein. The buffer was changed using desalting columns.

### Analytical ultracentrifugation

Sedimentation velocity experiments were performed using a Beckman Coulter Optima AUC and UV-Vis absorbance optics at 280 nm wavelength. An An-50 rotor and measuring cells with 12mm Epon centerpieces and quartz windows were used. Cleaning of measuring cells was performed by deionized water with 13.0 MΩ·cm resistivity, 20% ethanol and 1% Hellmanex detergent. All measurements were carried out at 50 000 rpm. Measurements were made in phosphate-buffered saline (137 mM NaCl, 2.7 mM KCl, 10mM Na_2_HPO_4_, and 1.8 mM KH_2_PO_4_, pH 7.35). CBM samples were measured in the concentration range from 0.05 g/L to 0.7 g/L that corresponds to 0.1 – 1.3 OD range. 400 μL of sample was loaded in each measuring cell. All experiments were performed at 20 °C and prior to each experiment the system was thermally stabilized for 1 h 30 min. To get complete sedimentation and sufficient amounts of data points, 600 scans per each sample were collected with a frequency of one scan per 80 seconds.

### Data analysis by Ultrascan

Data analysis were performed using the software UltraScan III (v. 4.0 revision 2466) (22). Rotor-stretching calibration and chromatic aberration corrections were applied during data import. A partial specific volume 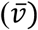 of 0.7148 mL/g was used. It was based on the amino acid sequence and calculations by the estimation tool implemented in Ultrascan. A theoretical density of 1.0056 g/mL and viscosity of 1.0120 mPa·s for the buffer were calculated using Ultrascan. During data editing, the meniscus position and end cutoff distance were entered manually. The plateau position corresponding to a uniform concentration prior to acceleration and the baseline buffer absorbance were determined automatically.

Concentration profiles obtained in the sedimentation velocity experiments were treated using 2-dimensional spectrum analysis (2DSA) (23). We used 2DSA to perform primary analysis coupled with simultaneous time-invariant and radial-invariant noise reduction to obtain the frictional ratio (*f*/*f*_*0*_), and sedimentation coefficient (*s*) values. The grid parameters used were 1 to 10 for *s* and 1 to 4 for f/f_0_ with 64 grid points in each direction. The choice of parameters was based on time-derivate analysis, and was selected to cover the distribution of parameters for all species in solution. The 2DSA was done in three steps. First, the time-invariant noise was removed after which the meniscus was fitted. The meniscus fit range was 0.03 cm around the previously chosen position. In the meniscus fitting procedure, 10 positions were chosen and corresponding root mean square deviation (RMSD) values were approximated by second order polynomial to determine the position with lowest RMSD. The position with lowest RMSD was used for further analysis. The third step was an iterative 2DSA refinement consisting of repetitions with improved meniscus position and fitting of the noise profile. The number of iterations was 10, which was sufficient for convergence of the fit. The parameters of the grid were the same in all steps.

The 2DSA results and noise reduction profile obtained by the analysis described above were used for further analysis. A genetic algorithm (GA) and a genetic algorithm Monte Carlo (GAMC) (24) optimization were used to extract the main populations of frictional ratio and sedimentation coefficient combinations present in the data. The existence of a reversible self-association was demonstrated using a van Holde-Weischet (vHW) (25) analysis. The dissociation constant (*K*_*D*_) was determined by a discrete model genetic algorithm (DMGA). The specific parameters used in GA, GAMC, vHW and DMGA are provided in the supplemental information. Hydrodynamic parameters of monomer and dimer were estimated using the UltraScan Solution Modeler (US-SOMO) (26). Calculations were performed using the ZENO hydrodynamic computation algorithm (27–29) and the van der Waals overlap bead model (30).

### Molecular modelling

All-atom molecular dynamics (MD) simulations were carried out by using the Gromacs software (version 5.1.4) (31,32). We used the particle Mesh Ewald (PME) electrostatics calculation scheme (33), and a non-bonded interactions cut-off of 1 nm. The modeling was performed with the Amber03ws force field (34) and TIP4P2005 water model (35). Two systems were studied, one with a single CBM_CipA_ molecule and one with two CBM_CipA_ molecules in the simulation box. A cubic box of 8 nm × 8 nm × 8 nm in initial size was used for the single CBM_CipA_ system and for the two molecule-system, a box size of 12.5 nm × 12.5 nm × 12.5 nm was used (Fig 1).

**Fig. 1.**
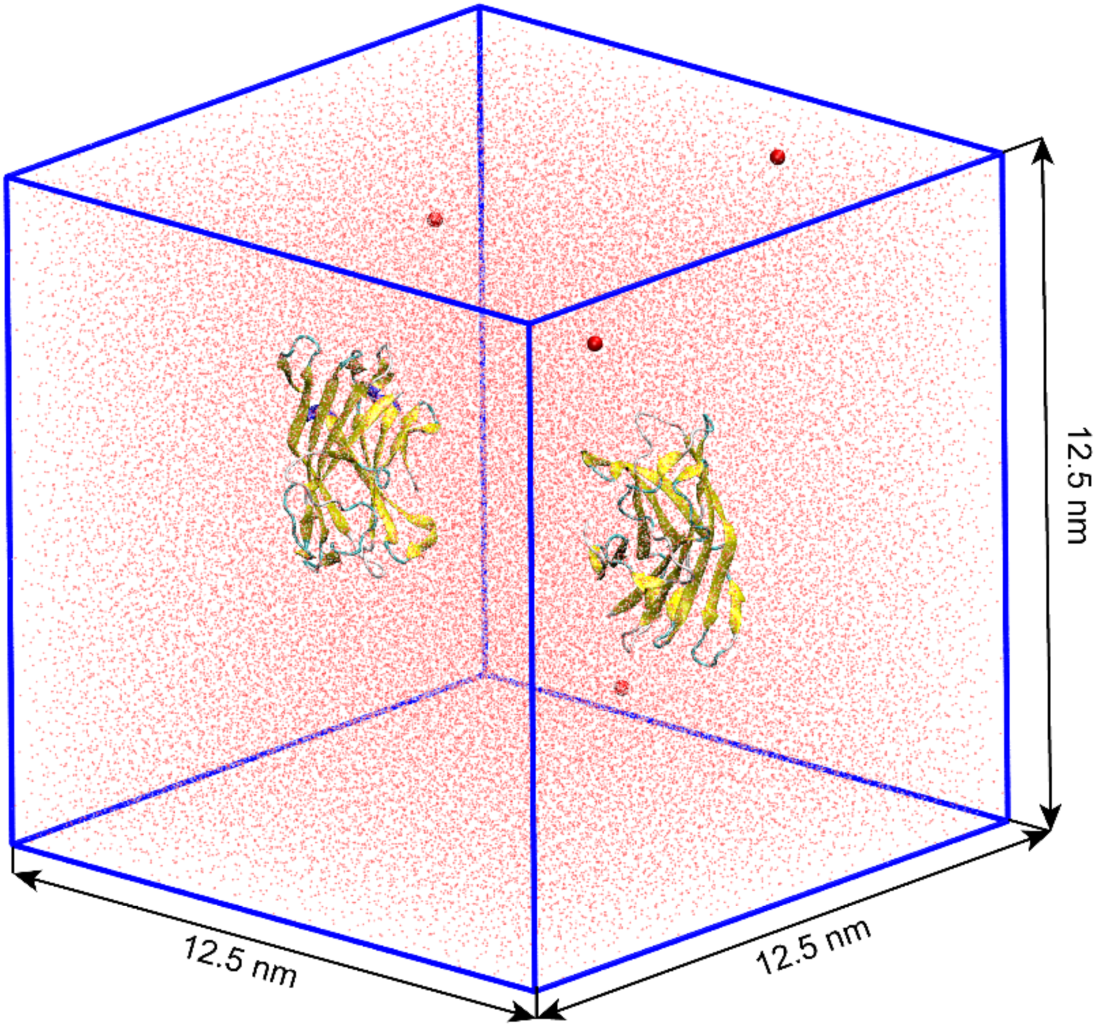
The initial all-atom molecular dynamics simulation configuration of two CBM_CipA_ molecules in the simulation box. The explicit water molecules are shown as red dots and the Na^+^ ions that are required for charge neutrality as red spheres.

The initial crystal structure of the single CBM_CipA_ simulations was the Protein Data Bank structure 1NBC. The file was modified to leave only the protein itself. The amino acid sequence differs from the sequence of the real protein by a missing polyHis-tag and two amino acids in the beginning and the end of the sequence. The initial structure for the two-molecule system was obtained using the final frame of the single CBM simulation, cutting the protein together with a 1 nm water shell. The initial arrangement of the two molecules relative to each other was chosen based on the location of the β-sheets and the geometric shape. After this, the proteins were solvated with explicit water molecules. Neutralization of the net charge in the system was performed by replacement of water molecules by Na^+^ ions: 4 ions were added in the two-molecule case. Both systems were energy minimized. After initial equilibration, the simulations were performed in the NPT (isothermal-isobaric) ensemble. The temperature and pressure control were carried out by the V-rescale thermostat (36) with a time constant of 0.1 ps and the Parrinello-Rahman barostat (37) with a time constant of 2 ps. The temperature and pressure reference levels were set at 300 K and 1 bar. Visualization was done with the VMD software (v. 1.9.4a9) (38). The main simulation was conducted for 160 ns for the single CBM_CipA_ and for 450 ns for the two-molecule system. In the two-molecule simulation, the first 150 ns were disregarded in the analysis.

Diffusion coefficients were calculated using the mean square displacement (MSD). The stable linear region on the MSD plot was approximated by a straight line according to the equation

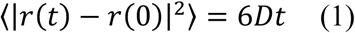

where ⟨|*r*(*t*) − *r*(0)|^2^⟩ is the MSD, *D* is the diffusion coefficient and *t* is the time.

## Results and Discussion

Sedimentation velocity experiments were conducted to investigate the possible oligomerization of CBM_CipA_ molecules. A set of solutions of CBM_CipA_ at different concentration were studied to follow changes in composition. After initial data treatment and noise reduction by 2DSA, GA was used further data refinement. An example of a sedimentation profile and fitted data by the GA at 0.7 g/L concentration is shown in Fig 2a. The GA analysis revealed two clearly identifiable peaks at 2.15 Sv and 2.7 Sv (Fig 2b, Table 1). These peaks correspond to molecular weights of 18 and 37 kDa. The 18 kDa peak at 2.15 Sv corresponds to the calculated molecular weight of CBM_CipA_ as a monomer which is 18469 Da. The peak at 2.7 Sv corresponds very closely to the mass of CBM_CipA_ as a dimer which is 36 938 Da. In addition, there was low intensity signal at 3.5 Sv that could represent minor amounts of higher oligomerization states but which is more likely to be an artefact due to random noise or sample impurities, as it did not show a concentration dependency. We conclude that the data show the presence of monomer-dimer system in which monomeric state prevails and is in good agreement of experimental and fitted data, and having a low overall level of noise.

**Table 1.** Sedimentation velocity fitting results for 7 different loading concentrations. Parameters of monomer and dimer peaks and also RMSD of the fitting are shown. Data for the dimer peak is shown only for the concentrations where the peak was clearly identifiable.

| Conc.<br>[mg/mL] | Monomer fraction |  |  |  | Dimer fraction |  |  |  |  |
| --- | --- | --- | --- | --- | --- | --- | --- | --- | --- |
| | $s$<br>[Sv] | $D \cdot 10^7$<br>[cm<sup>2</sup>/s] | $f/f_0$ | partial<br>conc. [%] | $s$<br>[Sv] | $D \cdot 10^7$<br>[cm<sup>2</sup>/s] | $f/f_0$ | partial<br>conc.<br>[%] | RMSD<br>of data<br>fitting |
| 0.05 | 2.09 | 9.8 | 1.26 | 95.4 |  |  |  |  | 0.00086 |
| 0.1 | 2.13 | 10.4 | 1.21 | 95.6 |  |  |  |  | 0.00102 |
| 0.2 | 2.15 | 10.4 | 1.19 | 91.4 |  |  |  |  | 0.00119 |
| 0.3 | 2.17 | 10.6 | 1.16 | 88.5 | 2.90 | 5.5 | 1.68 | 11.0 | 0.00144 |
| 0.5 | 2.19 | 11.0 | 1.16 | 82.9 | 2.76 | 5.3 | 1.74 | 12.6 | 0.00209 |
| 0.6 | 2.18 | 10.9 | 1.16 | 76.8 | 2.78 | 6.9 | 1.46 | 21.6 | 0.00248 |
| 0.7 | 2.19 | 10.9 | 1.16 | 74.7 | 2.81 | 6.8 | 1.47 | 24.0 | 0.00269 |
| Average | 2.16 | 10.5 | 1.2 |  | 2.8 | 6.0 | 1.6 |  |  |

**Fig. 2.**
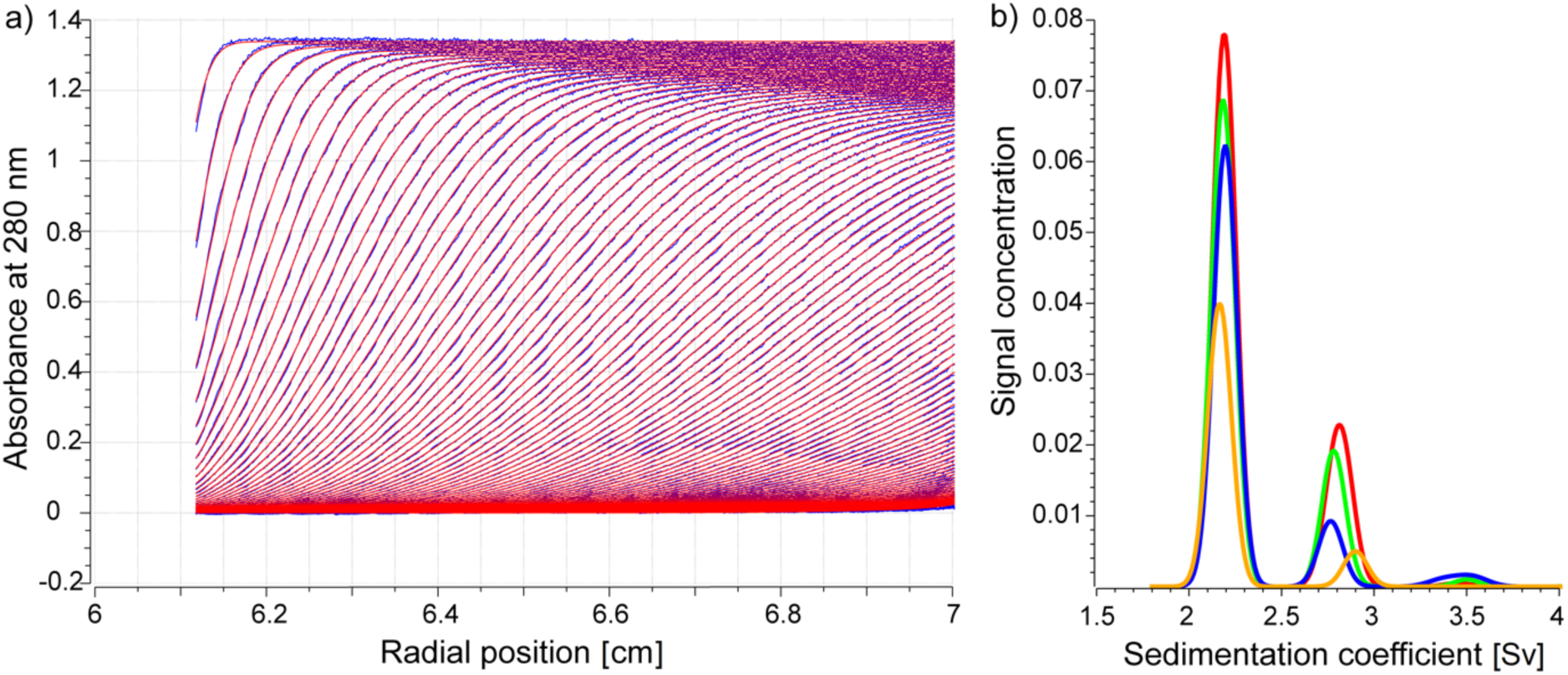
a) Sedimentation velocity experimental data (blue lines) and fitted curves based on the GA (red lines). b) GA analysis result shown as sedimentation distribution plot at four concentrations: 0.3 g/L – yellow, 0.5 g/L – blue, 0,6 g/L – green and 0,7 g/L red lines.

We conducted a GAMC analysis to evaluate the quality of data treatment and to perform confidence analysis by statistical methods. The pseudo 3Ds distribution connects sedimentation coefficients (*s*), frictional ratios (*f*/*f*_*0*_), and partial concentrations of the solution. The peak at 3.5Sv had a very low intensity and was not been taken for further analysis. No significant changes were observed in the pseudo 3D distribution after 64 Monte Carlo iterations (Fig 3). The analysis showed excellent stability and it allows us to conclude that both peaks were determined in correct way and definitely represent the solution composition.

**Fig. 3.**
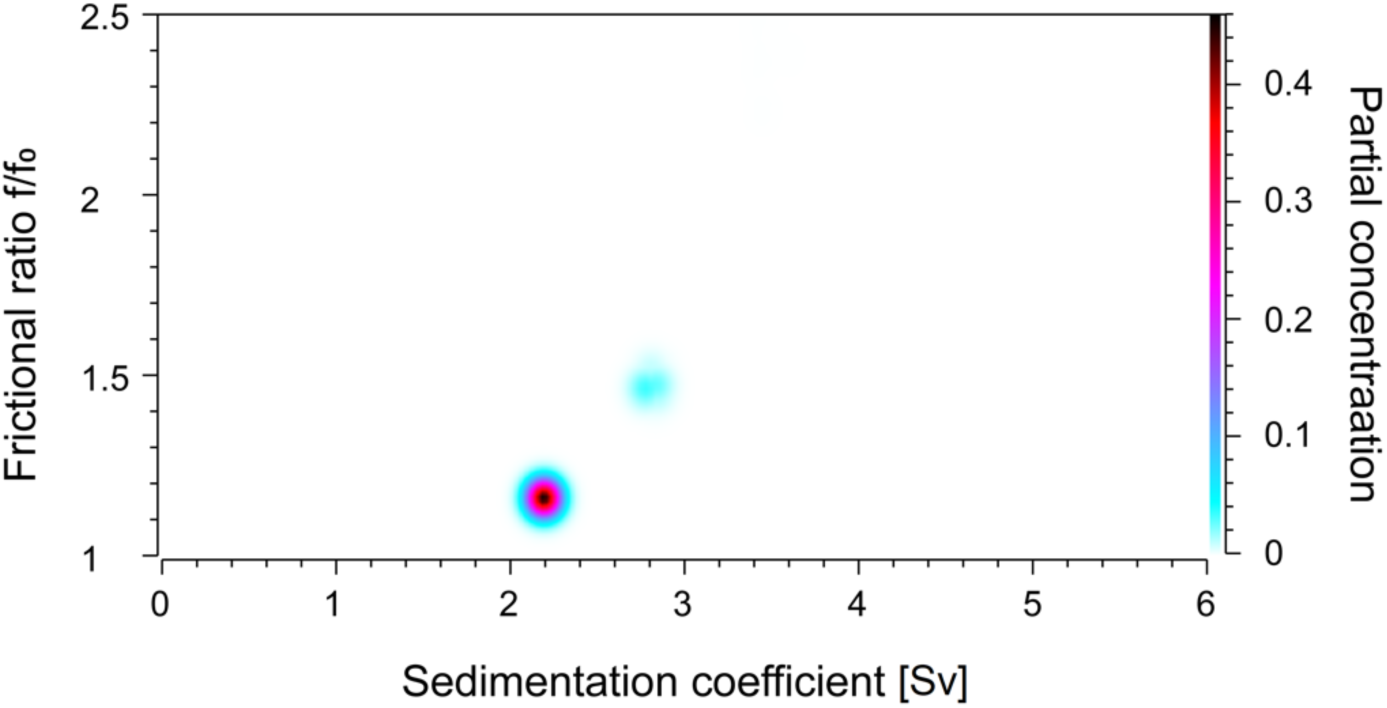
Pseudo 3D plot of the CBM_CipA_ solution composition at a concentration of 0.7 g/l after 64 Monte Carlo iterations by GAMC. The graph shows two peaks, a monomer at 2.2 Sv and f/f_0_≈1.15and a dimer at 2.8 Sv and f/f_0_≈1.5.

GAMC was then used to determine the diffusion constant (*D), s, f*/*f*_*0*_, and partial concentration values of the monomers and dimers for the full range of concentrations, i.e. from 0.05 to 0.7 g/l (Table 1). The sedimentation velocity fitting showed low RMSD values. The fitting was overall in good agreement with sedimentation data having only small changes in parameters at different concentrations. The CBM_CipA_ monomer showed an average *s* of 2.16 Sv and a *D* of 10.5·10^−7^ cm^2^/s. The *f*/*f*_*0*_ of the monomer was equal to 1.2 indicating that the monomer has compact globular structure as expected based on crystal structure. The dimer also showed expected values: a higher *s* value of 2.8 Sv, *D* = 6·10^−7^ cm^2^/s and a *f*/*f*_*0*_ of 1.6, representing a more anisotropic form. The partial concentrations of the two components showed a change with the total concentration, indicating the presence of dynamic interactions. In other words, the CBM_CipA_ behaved as reversible system. This is still further evidence that the peak at 2.7 Sv represents a dimeric form.

We used the vHW analysis as an alternative way to the GAMC analysis to investigate the occurrence of reversible oligomerization. The vHW method is based on a graphical transformation of the sedimentation velocity experimental data and allows obtaining a sedimentation coefficient distribution that is independent of diffusion. By the vHW method, one can identify boundary spreading due to diffusion and heterogeneity in the sedimentation coefficient as well as differentiation between non-interacting and self-associating species. If the kinetics are slow compared to the time of the experiment, the interacting components will be separated by the centrifugal force and the boundary of concentration profile will be separated. However, in the case of relatively fast kinetics the components will have time to re-equilibrate and a single boundary is observed. Changing the sample concentration will change the ratio between species causing changes in the boundary shape. This change represents a shift toward higher sedimentation coefficient and higher partial concentration of oligomers and can be seen in the vHW integral distribution. Such a shift in the vHW distribution at different concentrations was observed for CBM_CipA_ (Fig 4). The presence of the shift clearly indicates reversible self-association (21). Using DMGA we calculated the dissociation constant (*K*_*D*_) values for this reversible self-association (Table 2).

**Table 2.** The value of the dissociation constant K_D_ as determined at different concentrations.

| Conc. [mg/mL] ([μM]) | $K_D$, [μM] | 95% confidence interval | |
| --- | --- | --- | --- |
|  |  | lower | upper |
| 0.20 (10.83) | 81.3 | 68.8 | 93.8 |
| 0.30 (16.24) | 86.7 | 76.9 | 94.5 |
| 0.50 (27.07) | 93.9 | 81.3 | 106.6 |
| 0.60 (32.49) | 92.3 | 83.4 | 101.1 |
| 0.70 (37.90) | 80.5 | 68.8 | 92.2 |
| Average | 87.0 |  |  |

**Fig. 4.**
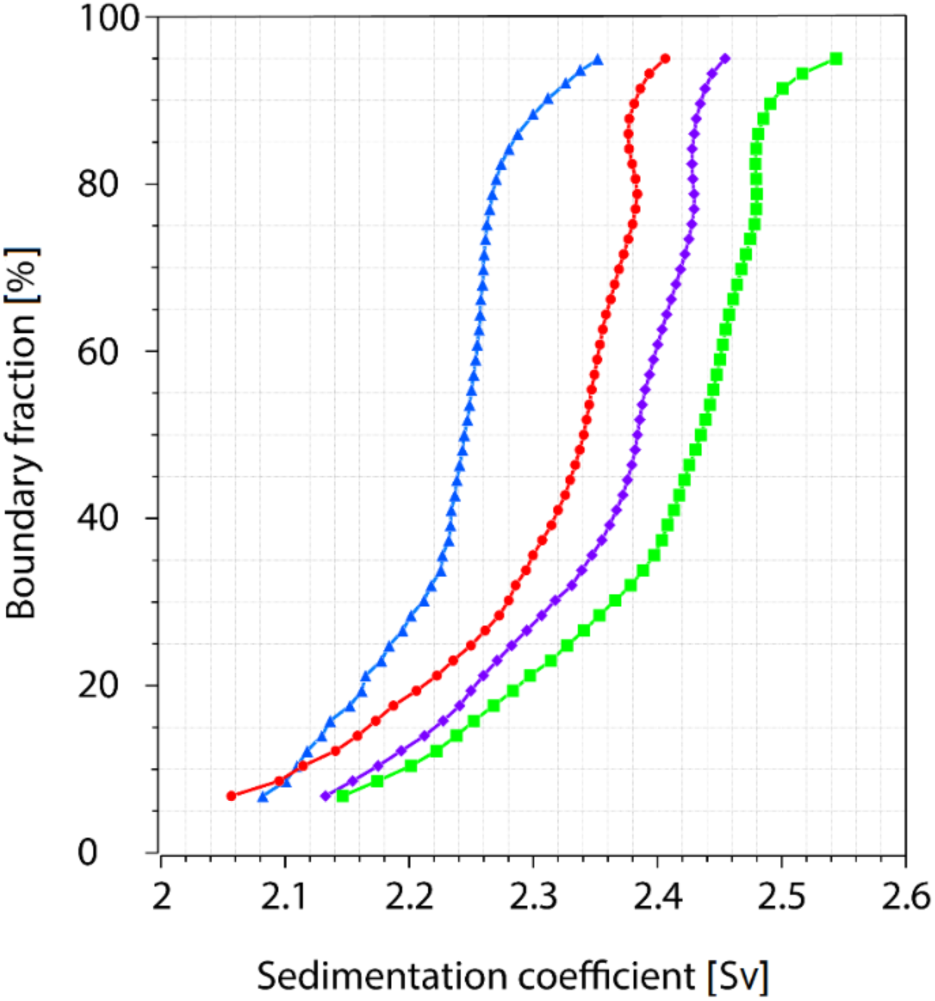
A van Holde-Weischet integral distribution plot for four different loading concentrations: 0.3 g/L – blue triangles, 0.5 g/L – red circles, 0.6 g/L – violet diamonds and 0.7 g/L – green squares. A clear shift of the curves towards higher Sv values correspond to a reversible self-association of the components.

Independently from the sedimentation velocity experiments, molecular dynamics simulations were also conducted. Two types of systems were simulated: one with a single CBM_CipA_ molecule and one system with two CBM_CipA_ molecules in the box. The first system was used to estimate the hydrodynamic properties of the CBM_CipA_ molecule as a monomer. From this, a diffusion coefficient of 7.9·10^−7^ cm^2^/s was calculated. Compared to the average measured *D* of 10.5·10^−7^ cm^2^/s, we consider the agreement good considering the widely different approaches used. One should note, however, that the MD simulations of the single CBM_CipA_ refer to the dilute solution in experiments as protein-protein interactions in the single CBM system would be interactions with periodic images via the periodic boundary conditions. Indeed, the monomer *D* decreases with the concentration (Table 1) and at the lowest concentration, i.e., 0.05 g/L, the *D* agrees slightly better with the MD results.

The second system allowed us to study the process of CBM dimerization. Initially, the CBMs were apart in the simulation box (Fig 1) but a stable dimer form was observed after 150 ns of MD simulations and the dimer form stayed the same until the end of simulation, i.e. next 300 ns (Fig 5). The CBM_CipA_ dimer diffusion coefficient was determined to be 6.2·10^−7^ cm^2^/s, which is in very good agreement with the measured one of 6.0·10^−7^ cm^2^/s. While the single monomer in the simulation box corresponds to concentration 6.0 g/L, the two CBMs in a simulation box correspond to concentration 3.1 g/L. The latter concentration is significantly above the concentration range studied in the experiments. This suggests, that the *D* determined from the MD simulations should actually be compared with the 0.7 g/L diffusion coefficient value, i.e. 6.8·10^−7^ cm^2^/s and not the dilute solution value.

**Fig. 5.**
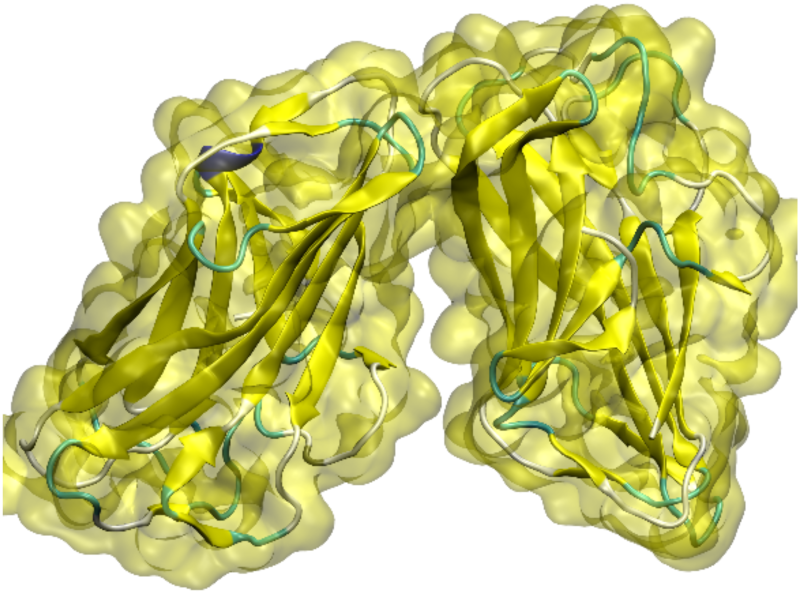
CBM_CipA_ dimer structure as obtained by the MD simulations. The snapshot corresponds to the final configuration at 450 ns.

For both the monomer and the dimer, the *D* calculated from the MD simulations is underestimated by 12-20 %. This might originate from the oversimplifications in the modelling methodology. For example, the *D* of the TIP4P/2005 water which is used in this work, is ∼10% smaller than experimental value (35). However, the MD method properly catches the decrease in *D* caused by dimerization.

A residue contact map (RCM) (Fig 6) was calculated from the two CBM simulation trajectory to understand which residues are involved in the dimer formation in the simulated system. The RCM shows the mean distance between the CBM residues, that is, each point on the map shows the distance between two residues, as specified by the residue index. We plotted the two CBM molecules with consecutive numbering, meaning that the lower left quadrant shows interactions within molecule 1, the upper right quadrant shows interactions within molecule 2, and both the lower right and upper left show interactions between molecules. Notably, an RCM is symmetric with respect to the diagonal from lower left to upper right. Residues on this line have a distance equal to zero, since each residue on this line is compared to itself. Square areas through which the zero-line passes display the distances between residues inside the molecule. The lines parallel and perpendicular to the zero line inside these areas identify parallel and anti-parallel β-sheets. The distance between residues belonging to different molecules is shown in square areas perpendicular to the zero line. The RCM revealed at least two areas which can participate in dimer formation and are involved in interactions between CBMs. The first area represents the interaction between residues Met-81 and Ser-82 of one CBM and Gly-65 of the other CBM. The second area shows interaction between Gln-48 of first molecule and group of residues Ser-63, Glu-102, Pro-103, Ala-105 belonging to the second molecule.

**Fig. 6.**
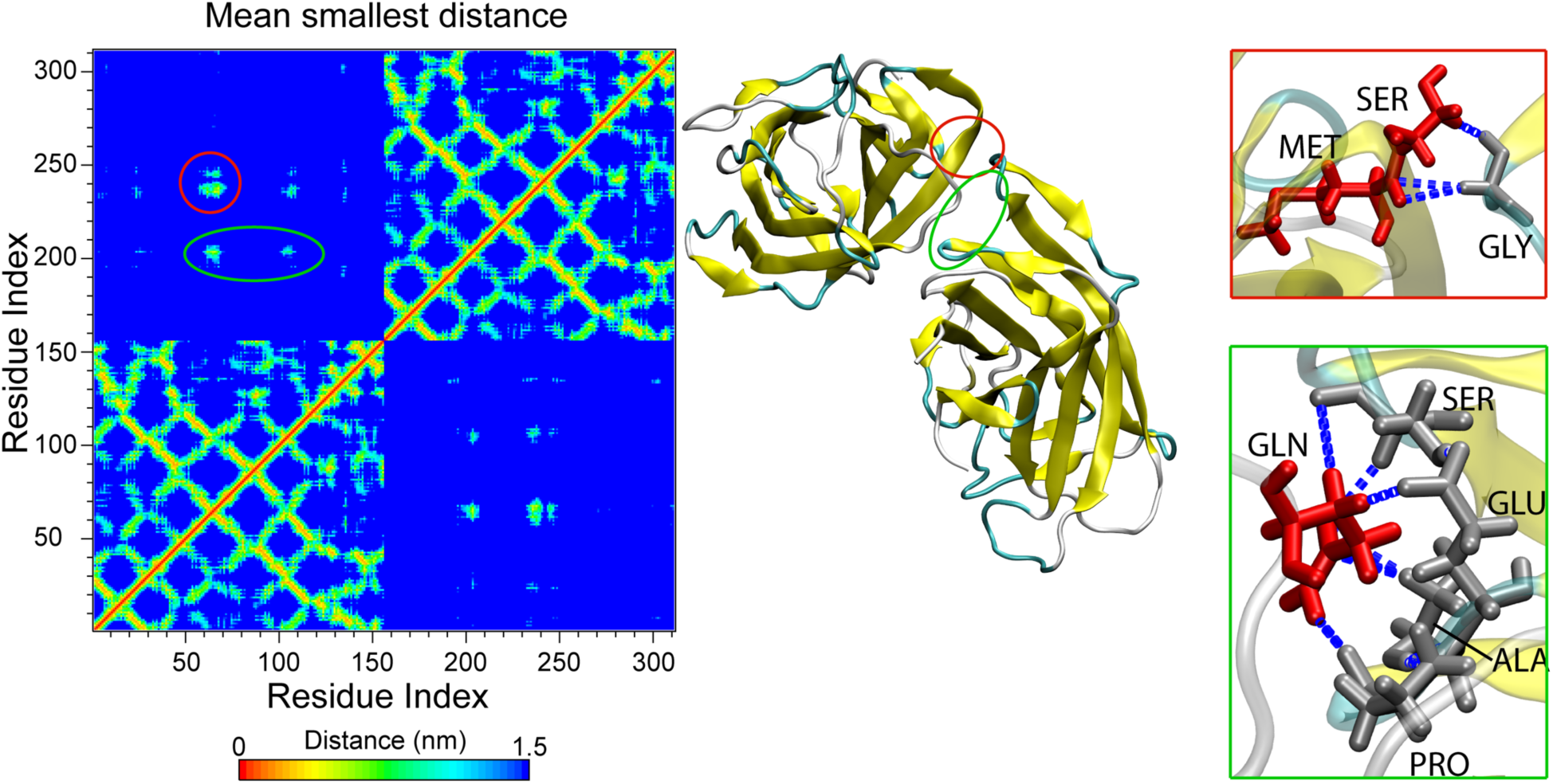
Residue contact map calculated for the two CBM molecular dynamics simulations and the corresponding dimer configuration with contact regions zoomed in. Red and green circles show areas of contact between molecules. Hydrogen bonds are shown by blue dot lines.

We also used SOMO to estimate the results of the sedimentation velocity experiments. The SOMO-estimations were based on the crystal structure for the monomer and the molecular dynamic simulations for the dimer. Table 3 shows that experimental and predicted values are in good agreement for the monomer but were slightly off for the dimer. Both sets of SOMO data show overestimation of *s* and *D* and underestimation of the *f*/*f*_*0*_. The slight difference between experimental and SOMO results could be explained by the absence of a poly-His tag and a few other amino acids in the crystal structure. In the case of the dimer, an inexact partial specific volume 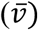 could also contribute.

**Table 3.**
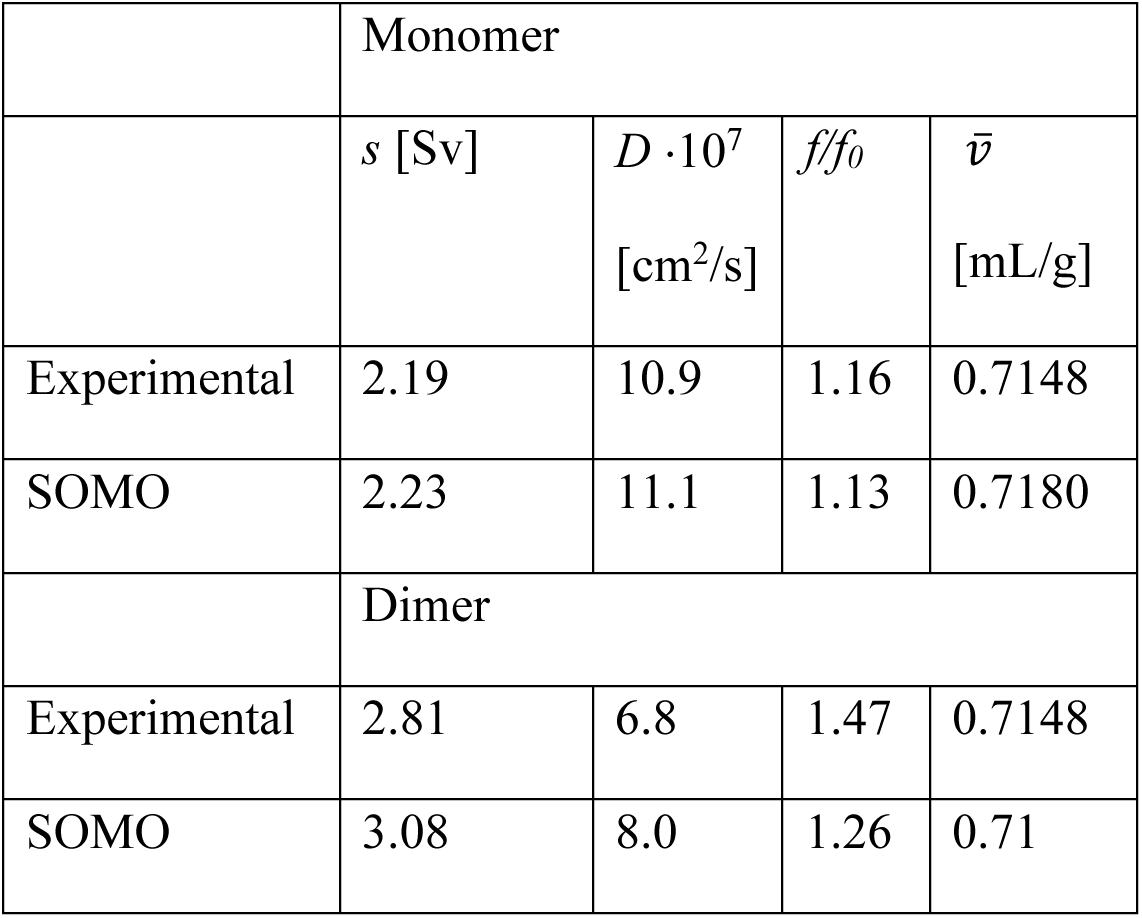
Comparison between experimental results at concentration 0.7 g/l and SOMO modelling.

## Conclusions

We showed that the CBM_CipA_ form dimers with a weak *K*_*D*_ of approximately 87 μM. The interaction can be classified as weak, on the edge of ultra-weak, as typically interactions that have a *K*_*D*_ over 100 μM are classified as ultra-weak (20). The *K*_*D*_ corresponds to a half of the CBM_CipA_ molecules being as dimers at a concentration of 1.6 g/L. As the *K*_*D*_ for the binding of CBM_CipA_ is at around 0.6 μM (13–15), which is around two orders of magnitude lower we conclude that its dimerization has a negligible effect on any function involving cellulose binding. However, in other uses of the CBM such as using then as part of molecular adhesives, much higher concentrations were used reaching over 2 mM (11,17). At these concentrations the majority of protein molecules are in the dimer form, unless the *K*_*D*_ is affected when the CBM_CipA_ is part of a fusion protein. Modelling indicated a possible mechanism of interaction between CBM molecules, and the modelled structures showed calculated properties such as diffusion and frictional ratio that had good agreement with the experimentally determined ones. In general AUC is suitable for studying weak and ultraweak interactions, and several examples are found. Approaches include both sedimentation velocity and sedimentation equilibrium setups and use of different software-packages for analysis (39–42). Meanwhile, the software capabilities are constantly developing and we found that the relatively recent approach of using the discrete model genetic algorithm (DMGA) method to determine the *K*_*D*_ to be a very efficient tool.

## Acknowledgements

We thank the Aalto School of Chemical Engineering graduate school for funding (DF). This work was supported by the Academy of Finland through its Centres of Excellence Programme (2014-2019, HYBER) and under Projects No. 307474 and 309324 (MS). We are grateful for the support by the FinnCERES Materials Bioeconomy Ecosystem and the Bioeconomy Infrastructure at Aalto. Computational resources by CSC IT Centre for Science and RAMI – RawMatTERS Finland Infrastructure are also gratefully acknowledged. Dr Sanni Voutilainen is acknowledged for cloning the CBM_CipA_. We thank professor Borries Demeler for his invaluable help with setting up Ultrascan.

## Supporting information

### Genetic algorithm (GA)

GA analysis is based on previously obtained 2DSA results. As an initialization step, bucket forms around each species. The width and the height of the bucket are automatically determined. We used autoassignment to define solute bins, however it is possible to assign bins and insert the parameters manually. Parameters of GA analysis were:

| Demes | Population size | Generations | Crossover percent | Mutation percent | Plague Percent | Elitism |
| --- | --- | --- | --- | --- | --- | --- |
| 1 | 200 | 100 | 50 | 50 | 4 | 2 |
| Migration rate | Regularization | Random seed | Simulation points | Band loading volume | Concentration threshold | S grid |
| 3 | 5 | 0 | 200 | 0.015 | 0.00001 | 100 |
| K grid | Mutate sigma | Mutate s | Mutate k | Mutate s and k |  |  |
| 100 | 0 | 20 | 20 | 20 |  |  |

### Genetic algorithm Monte Carlo (GAMC)

GAMC is based on the GA results. At this point we do the same steps as for GA and use the same parameters, only difference is that amount of Monte Carlo iterations equal to 64. We used parallel processing with 8 nodes to perform this type of analysis.

### van Holde-Weischet (vHW) analysis

van Holde-Weischet distribution plots were obtained by using the following parameters:

| Back diffusion tolerance | Divisions | Data smoothing | % of boundary | Boundary position [%] |
| --- | --- | --- | --- | --- |
| 0.201 | 50 | 30 | 90 | 5 |

In addition, about 10 – 15 first scans were deleted to decrease influence of instability in the beginning of the experiment on distribution.

### Discrete model genetic algorithm (DMGA)

The first step to perform DMGA is making the model and defining constrains.

The model consisted of two components: monomer and dimer, with the following parameters:

|  | Partial spec. volume [mL/g] | Extinction, OD [mol·cm] | Signal conc. | Molecular weight | Frictional ratio | Conc. dependency |
| --- | --- | --- | --- | --- | --- | --- |
| Monomer | 0.7148 | 35410 | 1.3436 | 18470 | 1.25 | 0 |
| Dimer | 0.7148 | 70820 | 0 | 36940 | 1.40 | 0 |

The signal concentration is the total concentration taken from final step of 2DSA.

DMGA Parameter constrains also were defined for monomer and dimer as follows:

The signal concentration is the total concentration taken from final step of 2DSA. DMGA Parameter constrains also were defined for monomer and dimer as follows:
| Demes | Population size | Generations | Crossover percent | Mutation percent | Plague Percent | Elitism |
| --- | --- | --- | --- | --- | --- | --- |
| 1 | 200 | 100 | 50 | 50 | 4 | 2 |
| Migration rate | Random seed | Simulation points | Band loading volume | Constraints parameter grid | Mutate sigma |  |
| 3 | 0 | 200 | 0.015 | 1000 | 0 |  |

